# Native Top-Down Proteomics of Endogenous Protein Complexes Enabled by Online Two-Dimensional Liquid Chromatography

**DOI:** 10.1101/2025.03.28.645965

**Authors:** Matthew S. Fischer, Holden T. Rogers, Emily A. Chapman, Song Jin, Ying Ge

## Abstract

Protein complexes are essential for virtually all biological processes, yet their structural characterization remains a major challenge due to their heterogeneous, dynamic nature and the complexity of the proteome. Native top-down mass spectrometry (nTDMS) has emerged as a powerful tool for comprehensive structural characterization of purified protein complexes, but its application to endogenous protein complexes in the proteome is challenging and typically requires labor-intensive and time-consuming prefractionation. Here, for the first time, we develop a nondenaturing online two-dimensional liquid chromatography (2D-LC) method for native top-down proteomics (nTDP), enabling high-throughput structural analysis of endogenous protein complexes. The automated, online interfacing of size-exclusion and mixed-bed ion-exchange chromatography achieves high coverage of endogenous protein complexes. We further develop a multistage nTDMS approach that enables comprehensive structural characterization within the chromatographic timescale, capturing intact non-covalent complexes, released subunits/cofactors, and backbone fragments. Our analysis detected 133 native proteoforms and endogenous protein complexes (up to 350 kDa) from human heart tissue in less than two hours. Such technological leaps in high-throughput structural characterization of endogenous protein complexes will advance large-scale nTDP studies in health and disease.

## Introduction

Protein complexes play essential roles in virtually all biological processes, with their functions governed by various structural factors including non-covalent interactions, post-translational modifications (PTMs), and genetic mutations.^1,2^ Structural characterization of protein complexes is crucial for understanding their dynamics and regulatory mechanisms in health and disease.^1–3^ Recently, native top-down mass spectrometry (nTDMS), which integrates native mass spectrometry^4–7^ with top-down proteomics,^8,9^ has emerged as a powerful tool for structural analysis of protein complexes.^10–14^ In nTDMS, proteins and protein complexes are ionized under non-nondenaturing conditions to preserve their high-order structure, non-covalent interactions, and PTMs, followed by gas-phase activation and tandem mass spectrometry (MS/MS) fragmentation to determine sequence, PTMs, and non-covalent binding sites.^10,12,13^ Thus nTDMS not only provides structural insights of macromolecular complexes, but also enables comprehensive analysis of proteoforms^15^—diverse protein variants resulting from alternative splicing, genetic variations, and PTMs.^10,12,16–18^ Despite its potential, nTDMS has been predominantly applied to purified protein complexes or high-abundance proteins.^10,13,16,19–21^

The high complexity of the proteome poses a major challenge for applying nTDMS to structurally characterize endogenous protein complexes at the proteome level, referred to as native top-down proteomics (nTDP), which requires the effective separation of intact proteins/complexes under non-denaturing conditions before MS analysis.^6,22,23^ The few reports that have successfully performed nTDMS for endogenous protein complex characterization at the proteome scale underscore the challenges.^11,24^ The initial large-scale nTDMS characterization of endogenous protein complexes relied on extremely labor-intensive and time-consuming offline prefractionation followed by direct infusion of hundreds of fractions into a mass spectrometer.^11^ Another study used offline size-exclusion chromatography (SEC) fractionation followed by native capillary zone electrophoresis (nCZE), but only identified smaller proteins complexes (<30 kDa).^24^ Such offline methods are time-consuming, prone to sample loss, and incompatible with high-throughput nTDP. Other recent efforts utilizing online one-dimensional (1D) separations such as nCZE^25^ or ion-exchange chromatography (IEC)^26^ were able to detect larger proteins but lacked MS/MS for identification of proteoforms and protein complexes.

To achieve both high-throughput and high-coverage structural analyses of endogenous protein complexes, herein we develop an online two-dimensional liquid chromatography (2D-LC) method interfacing nondenaturing SEC^27,28^ and IEC^26,29,30^ for nTDP (**Figure 1**). The separation efficiency is greatly increased in 2D-LC by leveraging a multiplicative relationship between the first and second separation peak capacities, or the theoretical number of resolvable components.^31,32^ Previously, online 2D-LC has been limited due to the mismatch of mobile phases between chromatographic dimensions. Therefore, here we implement active solvent modulation (ASM),^33^ a valve-based strategy that allows rapid transfer and dilution of MS-compatible mobile phase (i.e. ammonium acetate) between dimensions. Rapid automated transfer and modulation of effluent between chromatographic dimensions allows effective, orthogonal separation of protein complexes by charge and size prior to nTDMS structural analysis.

**Figure 1.**
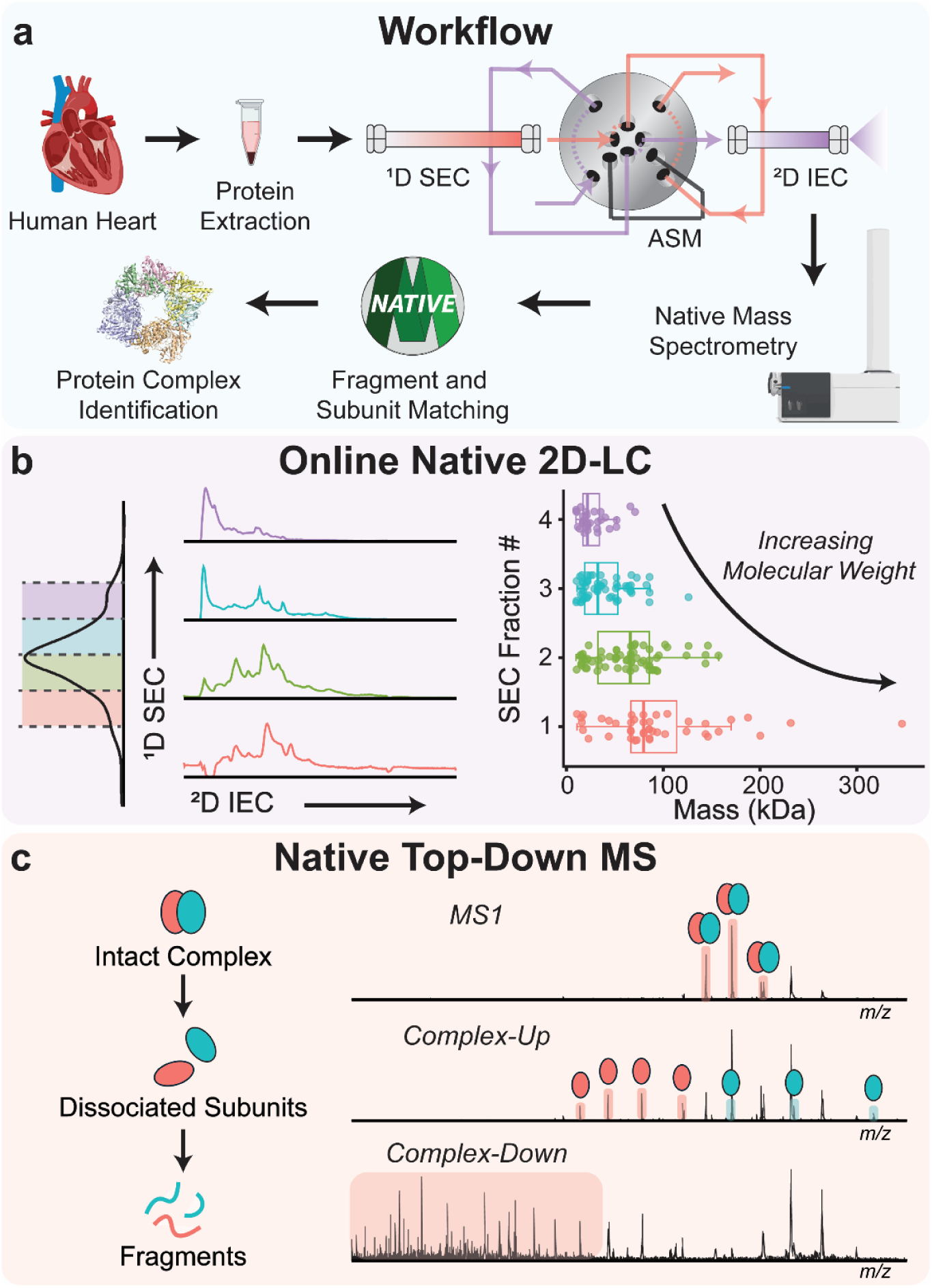
Native top-down proteomics platform featuring an online, MS-compatible multidimensional separation. **a)** Soluble proteins were extracted from human heart tissue and separated by nondenaturing online two-dimensional liquid chromatography (2D-LC) with active solvent modulation (ASM). The separation was coupled with a quadrupole time-of-flight (TOF) mass spectrometer for native top-down analysis. Identification was performed by matching intact complex, released subunit, and fragment masses with candidate proteins. The example structure is the creatine kinase S-type octamer. PDB: 4Z9M. **b)** Representative UV absorbance trace from ^1^D size-exclusion chromatography and corresponding total ion chromatograms (TICs) from the ^2^D mixed-bed ion-exchange chromatography (IEC) separations of each SEC fraction (*left*). Distribution of protein molecular weights detected in each SEC fraction (*right*). **c)** Endogenous protein complexes were subjected to native top-down MS analysis resulting in detection of the intact complex, released subunits (complex-up), and backbone fragment ions (complex-down).

We show that this online 2D-LC method greatly expands the proteome coverage achievable in nTDP with high throughput and automation. In a single automated 2D-LC analysis (<2 h), we have detected 133 native proteoforms and protein complexes, including large proteins up to 350 kDa, directly from human heart tissue. This technological advance will position nTDP as a critical tool for proteome-wide investigation into endogenous protein complex dynamics and function in health and disease.

## Results and Discussion

### Development and Characterization of Online Native 2D-LC

We select SEC-IEC as the optimal configuration for an online 2D-LC platform for nTDP considering factors including chromatographic resolution, mobile phase compatibility, and throughput. A major concern in top-down proteomics is the ion suppression of larger proteins and protein complexes caused by the coelution of smaller species.^34–36^ Therefore, we select SEC as the first chromatographic dimension (^1^D) because it separates proteins based on size, and, as there are ideally no interactions with the stationary phase, provides no reconcentration of analyte.^37^ Selecting a retentive chromatographic mode in the second dimension (^2^D) allows the dilute SEC effluent to be reconcentrated prior to detection, reducing peak width. Additionally, the lower resolution of SEC allows less frequent sampling of the ^1^D effluent, increasing throughput. This relationship can be visualized by the Davis Equation,^38,39^ which relates the overall peak capacity of a 2D separation to the ^1^D peak width and sampling time (**Figure S1**). IEC was selected as the ^2^D separation due to its high resolution, orthogonality, and good solvent compatibility with SEC (**Figure 1a, b**). Specifically, we selected mixed-bed IEC, which is capable of separating both basic and acidic proteins. We have previously demonstrated the ability of mixed-bed IEC to produce high-resolution, nondenaturing, MS-compatible protein separations.^30^

Nondenaturing protein SEC requires moderately high salt concentrations to avoid nonspecific interactions.^28,40,41^ Given the large ^2^D injection volumes used in online 2D-LC, significant analyte breakthrough occurred while transferring sample to the ^2^D IEC column. To understand this behavior, we determined the mixed-bed IEC retention characteristics of five standard proteins using the Linear Solvent Strength (LSS) model, a simple empirical model which can accurately predict protein elution in gradient separations.^42,43^ At the minimum ammonium acetate concentration we found necessary for SEC (200 mM), only the most strongly retained proteins would be trapped in IEC according to LSS calculations. We calculated that a salt concentration in the tens of millimolar was necessary for effective ^2^D IEC trapping and separation. ASM facilitated the 5× dilution of SEC effluent to approximately 40 mM prior to reinjection.^33^ Following ASM, a standard protein with low IEC retention was able to be retained and eluted by a salt gradient (**Figure S2, S3**).

To develop and characterize the SEC-IEC separation, we first used a mixture of four standard proteins of varying sizes and isoelectric points (**Table S1**). First, proteins were separated by 1D-LC in both SEC and IEC modes. In both cases, at least two standard proteins coeluted. Implementation of online 2D-LC successfully isolated all standard proteins to produce clean native mass spectra characterized by low charge states, narrow charge state envelopes, and preservation of non-covalent interactions (**Figure S4**). Following successful application to a standard protein mixture, we further developed the online SEC-IEC-MS/MS platform for nTDP analysis of a complex extract from human heart tissue (**Figure 1a**). This included optimization of the MS method to perform complex-up and complex-down native MS analysis for subunit dissociation and backbone fragmentation of endogenous protein complexes, respectively (**Figure 1c**).

### Native Top-Down Proteomics of Human Heart Protein Complexes

The heterogeneity of endogenous protein complexes presents a challenge to nTDP, especially when operating on the timescale of an online separation. In their native state, protein complexes have diverse structural properties which demand a wide range of MS conditions for desolvation, subunit dissociation, and backbone fragmentation.^10^ When analyzing a complex biological sample, it is difficult or impossible to develop a universal set of MS parameters to structurally characterize all protein complexes present. For example, a source energy which produces good desolvation of one complex may be too harsh and denature another smaller or more labile complex. To mitigate this problem, we designed a multistep MS method which varies source conditions and collision cell energy between SEC fractions (**Figure S5, Table S2**). This ensured that higher molecular weight protein complexes could be properly desolvated without denaturing lower molecular weight proteins. We also implemented alternating full-window collisionally activated dissociation (CAD) to cover a wide range of fragmentation energies for subunit dissociation and backbone fragmentation of diverse protein complexes. Three separate injections were performed with varied CAD energies in the alternating “MS/MS” scan to cover a wide range of protein complex labilities (**Table S2**). **Figure 2** provides an overview of the LC-MS workflow with representative data from two protein complexes, malate dehydrogenase (MDH, 72.7 kDa) and creatine kinase M-type (CKM, 85.9 kDa). Both proteins were detected as homodimers in the second SEC fraction, with low charge states, narrow charge envelopes, and average mass accuracy within 1 Da (**Figure 2a, b**). Upon collisional activation, monomers were released from each complex (**Figure 2c**). Activation of MDH released subunits with a wide range of charge states: we detected charge-stripped monomers with charge states as low as +5. At higher CAD energies, backbone fragments were detected and matched to the sequences of MDH and CKM (**Figure 2d**).

**Figure 2.**
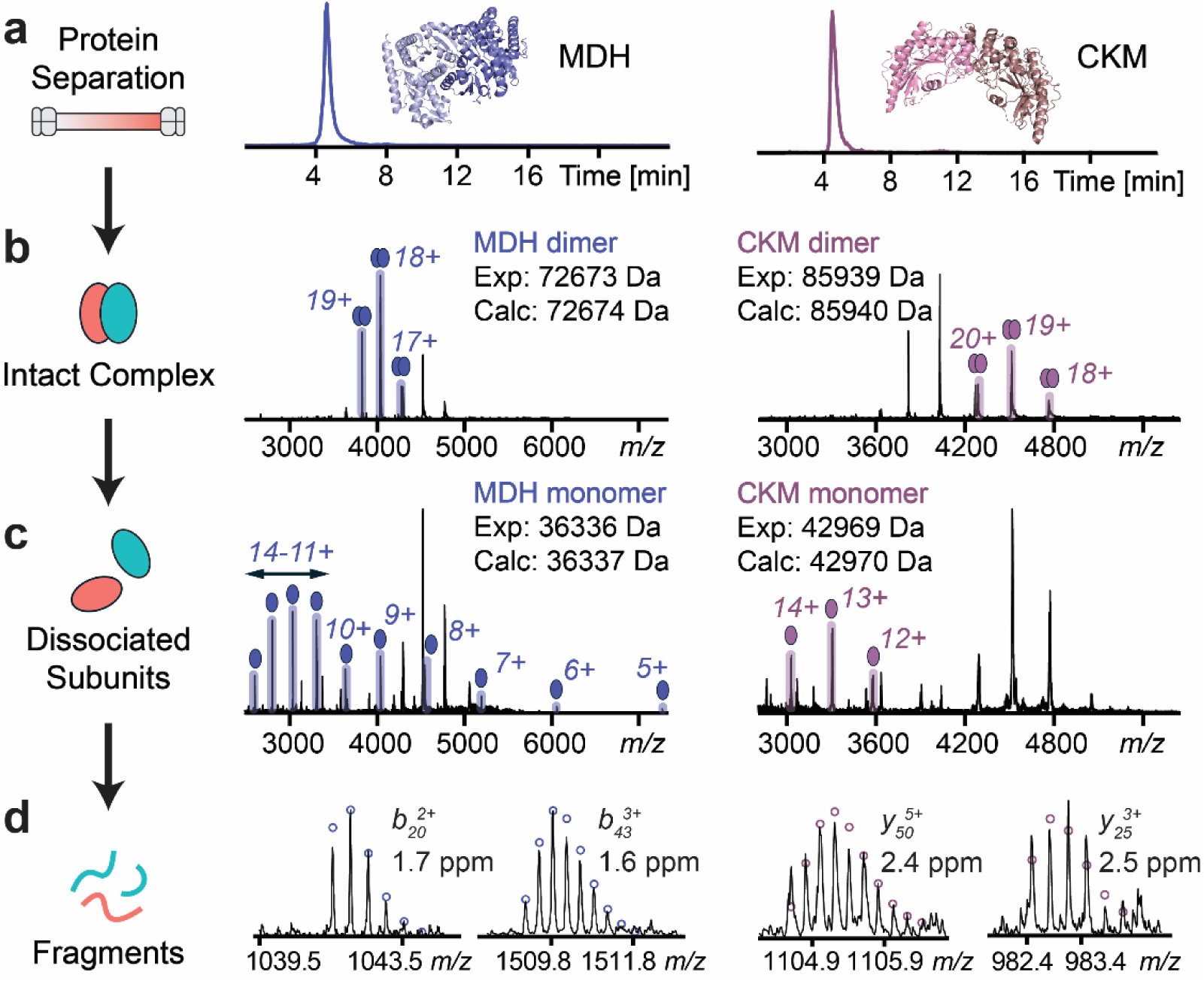
Representative native top-down MS analysis of endogenous protein complexes from human heart tissue. **a)** Extracted ion chromatograms (EICs) for malate dehydrogenase (MDH) and creatine kinase M-type (CKM). EICs were generated from the ion-exchange separation of the second fraction of the ^1^D size-exclusion separation. PDBs: 7RM9 and 1I0E. **b)** Detection of the intact MDH and CKM complexes. **c)** Detection of released subunits from MDH and CKM upon raising the collisional energy during the alternating “MS2” scan phase. **d)** Representative fragment ions from both proteins with theoretical isotopic fits and high mass accuracy.

A full 2D-LC experiment involved transferring four fractions from ^1^D SEC to ^2^D IEC. In the first SEC fraction, corresponding to the largest proteins, we successfully dissociated subunits from a set of tetrameric glycolytic enzymes (**Figure 3**). This included glyceraldehyde 3-phosphate dehydrogenase (G3P, 144.7 kDa), L-lactate dehydrogenase B (LDHB, 146.2 kDa), fructose-bisphosphate aldolase A (ALDOA, 157.2 kDa), and pyruvate kinase isoform M1 (PKM1, 231.9 kDa). For each tetramer, increasing collisional activation resulted in the dissociation of highly charged monomers, each detected within 1 Da mass accuracy (**Figure 3c**). A few backbone fragment ions were detected at the tested CAD energies (**Figure S6**).

**Figure 3.**
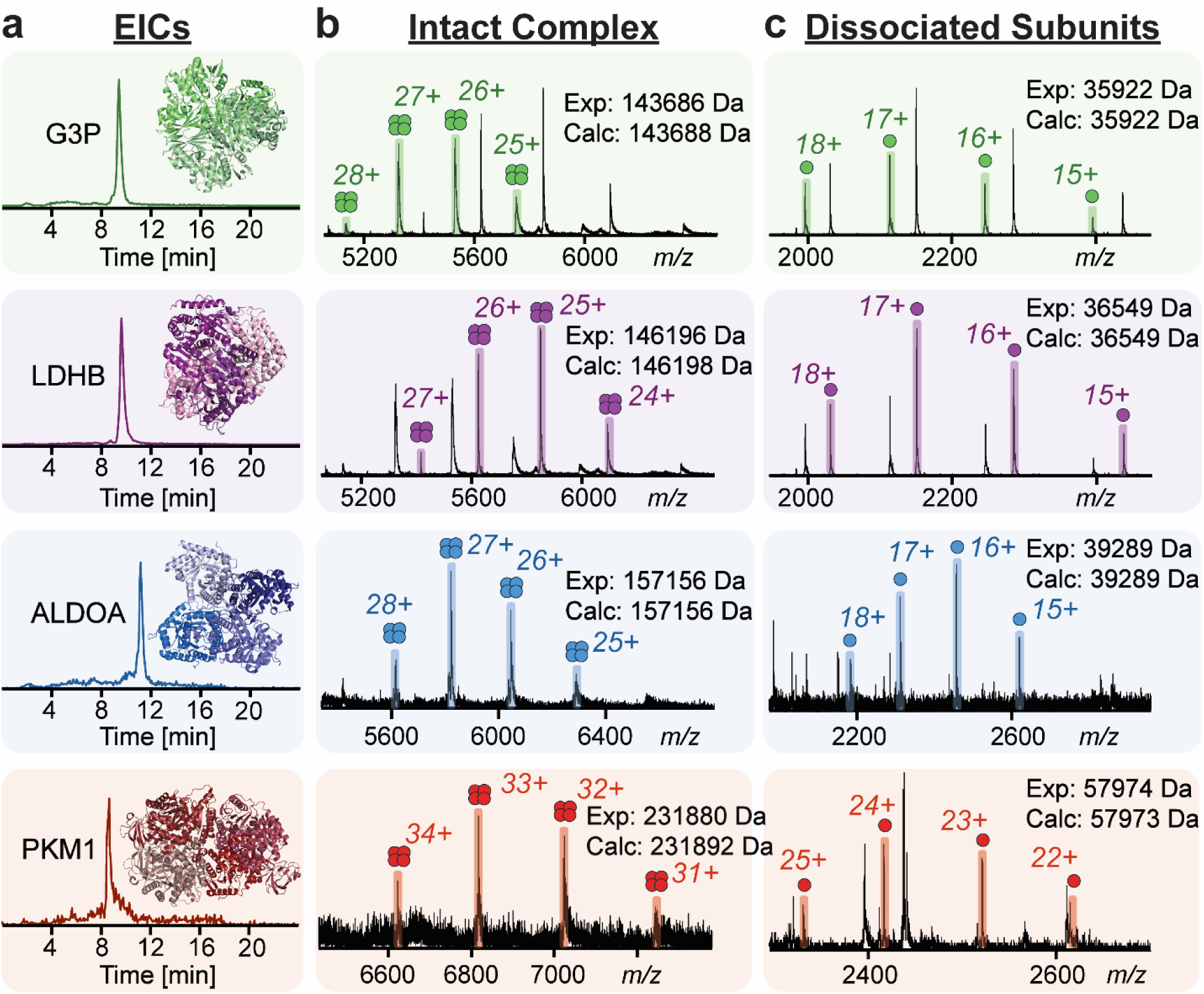
Representative complex-up analyses of tetrameric glycolytic enzymes. **a)** Extracted ion chromatograms (EICs) for glyceraldehyde 3-phosphate dehydrogenase (G3P), L-lactate dehydrogenase B (LDHB), fructose-bisphosphate aldolase A (ALDOA), and pyruvate kinase isoform M1 (PKM1). EICs were generated from the ion-exchange separation of the first fraction of the ^1^D size-exclusion separation. PDBs: 1U8F, 1T2F, 5KY6, and 3SRF. **b)** Detection of the intact complex of each enzyme. **c)** Complex-up experiment involving the dissociation of subunits from each tetramer upon collisional activation.

In some cases, either complex-up or complex-down data could not be obtained for a protein complex. For example, in one spectrum we detected mitochondrial succinyl-CoA:3-ketoacid coenzyme A transferase 1 (OXCT1, 104.2 kDa dimer). We were able to detect backbone fragments from OXCT1, but not released subunits (**Figure S7, S8**). Similar behavior has been reported in previous work studying higher-order structure of protein complexes with CAD.^16^ In the same elution window where OXCT1 was detected, we detected the 170 kDa mitochondrial enoyl-CoA hydratase (ECHM) hexamer and its dissociated subunits (**Figure S7**). Additionally, we detected the release of isochorismatase domain-containing protein 1 (ISOC1, 32.1 kDa) which, according to UniProt, is not known to form a complex with any other proteins (**Figure S7, 9**). Our nTDP method is also capable of detecting the binding and release of cofactors, shown by the detection and analysis of hemoglobin in the third SEC fraction (**Figure S10**). We successfully detected the intact heterotetrametric hemoglobin complex with four heme cofactors, followed by dissociation into alpha and beta subunits, release of heme, and backbone fragmentation of the subunits.

The reproducibility of the SEC-IEC method was assessed by triplicate analyses of human heart extract, and the coverage was compared to 1D-LC analyses of the same sample. Triplicate injections showed good reproducibility in both dimensions (**Figure S11**). Compared to 1D SEC and 1D IEC, online 2D-LC detected many more native proteoforms, especially high molecular weight species (**Figure S11, S13**). Specifically, 21 native proteoforms were detected by the 1D SEC approach, and 46 native proteoforms were detected by 1D IEX (**Table S3**). In contrast, 133 native proteoforms and protein complexes were detected from one online 2D-LC injection with a duration under two hours (**Table S4**). Of these, 36 native proteoforms including 27 protein complexes were identified by a combination of MS/MS strategies (**Table S5**). Additional examples of detected protein complexes can be viewed in **Figure S14**. Although this method is promising, there are areas which could be improved. The sensitivity of the method could be greatly increased (and sample consumption reduced) by scaling the separation down to microscale or nanoscale, though this would create new considerations and hardware requirements for online 2D-LC. Achieving efficient fragmentation of diverse protein complexes on the LC timescale is challenging, and our study was performed using an Agilent 6545XT Q-TOF mass spectrometer with only CAD fragmentation capabilities and a quadrupole isolation limit of 4000 *m/z*. The implementation of higher *m/z* isolation capabilities and expanded top-down fragmentation platforms (e.g. ExD cell^44^) would greatly complement online 2D-LC-enabled nTDP.

### Identification of a Large Protein Complex (>340 kDa)

In our SEC-IEC-MS/MS experiments, backbone fragmentation from complex-down experiments was not always sufficient for unambiguous protein identification, especially for larger or less abundant protein complexes. However, in many cases there was a likely candidate for the identity of a detected protein on the intact complex or dissociated subunit level based on previously published proteomics data from human heart extracts.^36,45^ To unambiguously identify these proteins, we performed an additional experiment where the ^2^D IEC separation was exchanged for RPLC (**Figure 4a**). In this configuration, proteins remained in their native state throughout the ^1^D separation, being subsequently trapped and denatured on the ^2^D RPLC column. Using RPLC as the second dimension provided high chromatographic resolution and efficient fragmentation, given the higher fragmentation efficiency of denatured versus native proteins.^46^

**Figure 4.**
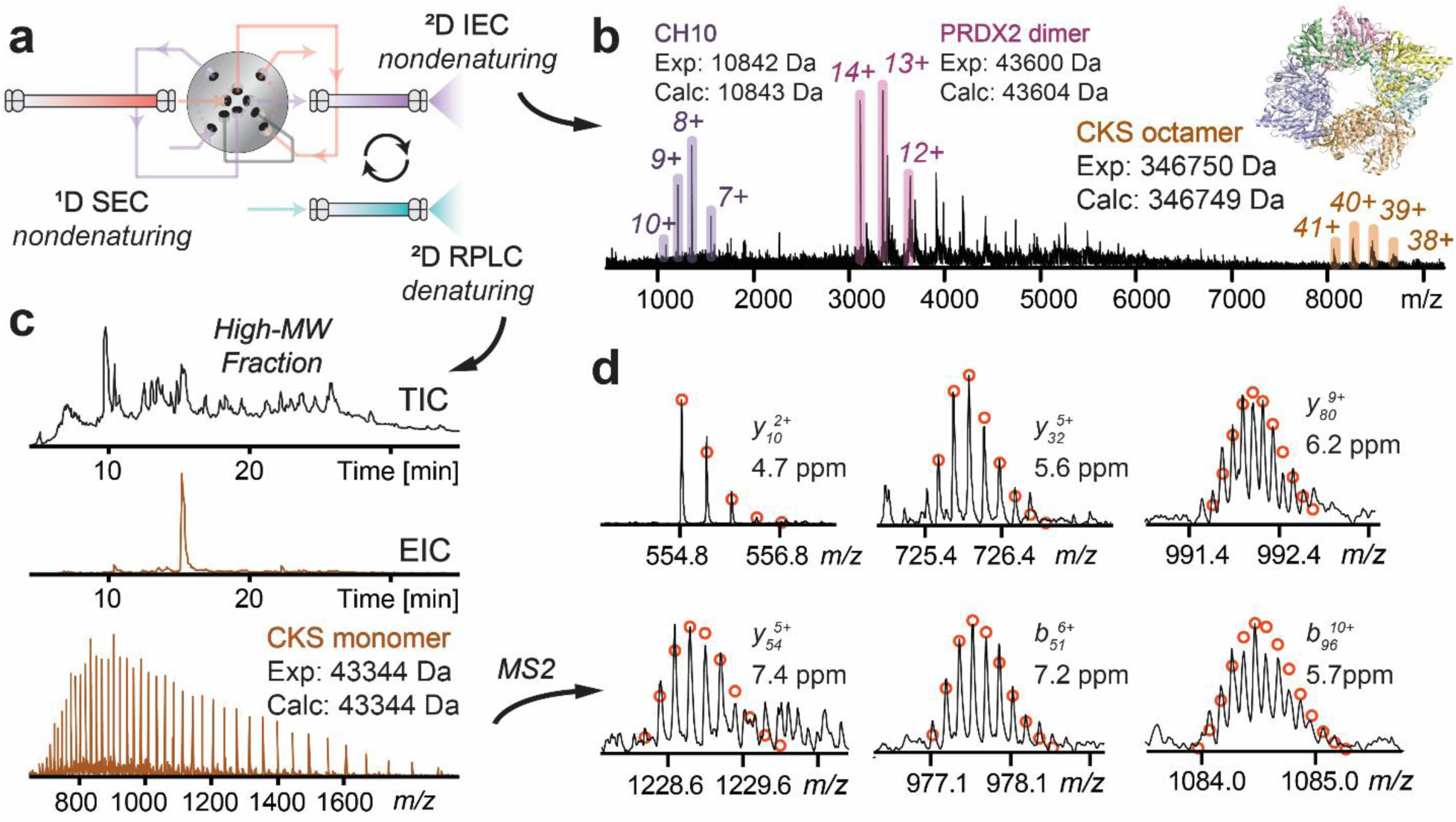
Detection and identification of the creatine kinase S-type octamer. **a)** Schematic describing the two online 2D-LC configurations used. To confirm the identity of CKS, the second dimension (^2^D) separation was exchanged for reversed-phase liquid chromatography (RPLC). **b)** Native mass spectrum of the creatine kinase S-type (CKS) octamer and two coeluting species: 10 kDa heat shock protein (CH10) and peroxiredoxin-2 (PRDX2). PDB: 4Z9M. **c)** Total ion chromatogram (TIC) from the RPLC separation of the first SEC fraction, where CKS was originally detected. Extracted ion chromatogram (EIC) and mass spectrum corresponding to the CKS monomer. **d)** Representative fragment ions and theoretical isotopic fits from collisionally activated dissociation (CAD) of the CKS monomer.

With SEC-RPLC-MS/MS, we were able to identify the largest protein we had previously detected as mitochondrial creatine kinase, S-type (CKS, 346.7 kDa octamer) (**Figure 4b**). In our native 2D-LC experiments, we detected the CKS octamer in the 8000-9000 *m/z* range, with 40+ as the most abundant charge state. Interestingly, we detected the mitochondrial 10 kDa heat shock protein (CH10) and disulfide-linked peroxiredoxin-2 (PRDX2) dimer in the same spectrum (**Figure 4b**); this was surprising given their low molecular weights. Both species are known to form much larger protein complexes: PRDX2 forms a ring-like decamer, and CH10 forms a homoheptamer which ultimately interacts with CH60 to form the football-like mitochondrial chaperonin complex.^47^ Given the presence of these proteins in the first SEC fraction, it is possible that these complexes are dissociating either during the IEC separation or during ionization and activation for mass spectrometry. Indeed, the denatured CH60 monomer was detected in this same SEC fraction by SEC-RPLC-MS/MS (**Figure S15**). Additional data and examples of protein complexes verified by SEC-RPLC-MS/MS are included in **Figures S16-S17**.

Here we have developed a novel nTDP method coupling online 2D-LC with nTDMS, to enable high-throughput structural analysis of endogenous protein complexes with high proteome coverage. In a single automated 2D-LC analysis (<2 h), our method detected 133 native proteoforms and endogenous protein complexes directly from human heart tissue, including large protein complexes up to 350 kDa. Notably, we detect large protein complexes, with an average protein mass of 94 kDa in the first SEC fraction and the detection of species up to 350 kDa. In comparison, the most comprehensive prior nTDP study identified 160 proteins (with 81 of these forming 125 proteoform complexes) from ∼600 fractions from mouse hearts and four human cancer cell lines, with each fraction being manually analyzed by direct infusion into a mass spectrometer, an extremely laborious and time-consuming process.^11^ Another study used offline SEC fractionation followed by online nCZE-MS, detecting 23 protein complexes under 30 kDa and an average proteoform mass of 13 kDa.^24^ Additionally, a recent online nCZE study using an ultra-high mass range Orbitrap detected species up to ∼400 kDa but lacked MS/MS for structural characterization and faced limitations in reproducibility and peak capacity (achieving an estimated peak capacity of 15).^25^ Our method achieves high-throughput and efficient separations (peak capacity = 70) of endogenous protein complexes while providing high proteome coverage and identification with top-down MS/MS analysis. Continued development of nTDP bioinformatics tools can help accelerate the identification of protein complexes.

The flexibility of online 2D-LC enables potential integration of different chromatographic modes such as HIC^48^ for orthogonal, nondenaturing separation, enhancing proteome coverage. Furthermore, sensitivity could be greatly improved by developing an equivalent nanoscale 2D-LC method. Integration with high *m/z* range quadrupole technology and ion activation methods such as ultraviolet photodissociation (UVPD)^49^, surface-induced dissociation (SID)^50^, and electron-based fragmentation methods (ExD)^51^ could further enhance protein identification and structural characterization.

## Conclusions

In this study, we developed an online 2D-LC-enabled nTDP platform for high-throughput structural characterization of endogenous protein complexes directly extracted from biological samples. The automated interfacing of MS-compatible, nondenaturing SEC and IEC provided efficient separations of protein complexes in their native states, including high molecular weight multimeric protein complexes. From a single experiment (< 2 h), we detected 133 native proteoforms and protein complexes (up to 350 kDa) from human heart tissue. We developed a multistage nTDMS method which allowed detection of released subunits and backbone fragment ions from a range of protein sizes and properties. With this method, we performed complex-up analysis of proteins including the 170 kDa ECHM hexamer and 232 kDa PKM1 tetramer. The flexibility of online 2D-LC also allowed facile integration of RPLC for unambiguous protein identification and proteoform characterization of unknown complexes. This study presents a major technological leap in nTDP for high-throughput structural characterization of endogenous protein complexes. Future advancements in the sensitivity of 2D-LC and gas-phase fragmentation strategies will further establish nTDP into an indispensable tool for proteome-wide exploration of the structure, dynamics, and function of endogenous protein complexes in health and disease.

## Experimental Section

### Reagents and Consumables

All reagents were purchased from MilliporeSigma (Burlington, MA, USA) unless otherwise noted. Solvents were prepared with water from an in-house Milli-Q system (Millipore, Corp., Billerica, MA, USA). Ion-exchange and size-exclusion columns were generously provided by PolyLC Inc. (Columbia, MD, USA), and the reversed-phase column was purchased from Waters Corp. (Milford, MD, USA).

### Human Cardiac Tissue Collection

Healthy donor hearts with no history of heart disease were obtained from the University of Wisconsin–Madison Organ and Tissue Donation-Surgical Recovery and Preservation Services. Tissue was stored in cardioplegic solution until dissection and then flash-frozen in liquid nitrogen and stored at −80 °C. The procedures for the collection of human donor heart tissues were approved by the UW–Madison Institutional Review Board.

### Protein Extraction from Human Heart Tissue

Human heart tissue (20 mg) was homogenized in 150 μL of HEPES extraction buffer (25 mM HEPES pH=7.4, 60 mM NaF, 1 mM L-methionine, 1x HALT protease/phosphatase inhibitor cocktail) using a Teflon pestle at 4 °C. The homogenate was centrifuged for 30 min at 21,000 × *g* at 4 °C and the supernatant was transferred to Eppendorf Protein Lo-Bind tubes, snap frozen in liquid nitrogen, and stored at −80 °C. Prior to analysis, protein extracts were thawed on ice and desalted into the starting mobile phase composition using an Amicon 30 kDa molecular weight cut-off filter. A Bradford protein assay was used to determine the final concentration.

### 1D-LC and 2D-LC Analyses of Standard Proteins and Human Heart Extracts

All liquid chromatography–mass spectrometry (LC-MS) experiments were conducted with an Agilent 1290 Infinity II 2D-LC System and an Agilent 6545XT AdvanceBio LC/Q-TOF with an Agilent Jet Stream electrospray ionization source. LC experiments without MS were performed on either the Agilent 1290 system or a Waters ACQUITY UPLC H-Class with detection by UV absorbance at 280 nm. A PolyLC CATWAX column (2.1×100 mm, 3 μm, 1500 Å) was used for all IEC experiments, and a PolyLC HYDROXYETHYL A column (2.1×200 mm, 3 μm, 500 Å) was used for all SEC experiments.

For LC-MS development, the standard protein mixture contained enolase (Eno; 93 kDa), alcohol dehydrogenase (ADH; 147 kDa), ovalbumin (Ova; 44 kDa), and trypsinogen (Trp; 24 kDa) at approximate concentrations of 5.0, 10.0, 5.0, and 0.5 mg/mL, respectively. The complete list of standard proteins used for LC-MS development and Linear Solvent Strength modeling is included as **Table S1**, along with their accession numbers. The mixture was exchanged into 10 mM ammonium acetate using an Amicon 10 kDa molecular weight cut-off filter. For the SEC separation, the mobile phase was 200 mM ammonium acetate, and the flow rate was 30 μL/min. For the IEC separation, the mobile phases were 10 mM ammonium acetate (A) and 500 mM ammonium acetate (B) with a gradient of 0-100-0-0 %B in 0-15-15.01-25 minutes and a flow rate of 200 μL/min. For the 2D-LC experiment, the sample transfer loops had a volume of 40 uL, and the ASM factor was 5.1. The ^2^D IEC gradient was 0-100-0-0 %B in 0-10-10.01-18 minutes. The columns were kept at room temperature.

For the 1D SEC analysis of human heart protein extract, the mobile phase was 200 mM ammonium acetate with a flow rate of 30 μL/min. For the 1D IEC analysis of human heart protein extract, the mobile phases were 10 mM ammonium acetate (A) and 500 mM ammonium acetate (B) with a gradient of 0-100-0-0 %B in 0-30-30.01-40 minutes and a flow rate of 200 μL/min. Flow was split 3:1, resulting in 50 μL/min to the MS.

For the 2D-LC analysis of human heart extracts, flow exiting the IEC column was split 3:1, resulting in 50 μL/min to the MS. The ^2^D IEC gradient was 0-100-0-0 %B in 0-15-15.01-23 minutes. To assess the reproducibility of the system, three replicates of 60 μg of protein were injected. For native proteomics, three additional injections of 120 μg of protein were conducted. For the SEC-RPLC analysis, the ^2^D IEC column was exchanged for a Waters BioResolve RP mAb Polyphenyl column (2.1×150 mm, 2.7 μm, 450 Å). Only the first SEC fraction was collected and subjected to RPLC. The RPLC mobile phases were 0.1% formic acid in water (A) and 0.1% formic acid in ACN (B) with a gradient of 5-5-30-60-95-95-5-5%B in 0-2-5-50-50.01-55-55.01-60 minutes. The flow rate was 200 μL/min, flow splitting was removed, and 60 μg of protein was injected.

### MS Method for Online 2D-LC Native Top-Down Proteomics

For native proteomics, a scan rate of 0.5 Hz was used, alternating between low and higher energy in the collision cell. The higher energy scan served as a full window MS/MS scan for subunit dissociation and fragmentation of native protein complexes. The source conditions and collisional energies were also varied between SEC fractions and between three separate injections to cover a wide range (**Table S2**). The source parameters which remained constant were the gas temperature of 150 °C, drying gas flow of 10 L/min, nebulizer pressure of 30 psi, sheath gas temperature of 100 °C, sheath gas flow of 10 L/min, capillary voltage of 5000 V, and nozzle voltage of 2000 V.

For the SEC-RPLC experiments, the source conditions were a gas temperature of 350 °C, drying gas flow of 12 L/min, nebulizer pressure of 30 psi, sheath gas temperature of 350 °C, sheath gas flow of 12 L/min, capillary voltage of 5000 V, and nozzle voltage of 2000 V. The fragmentor and skimmer voltages were set to 180 V and 65 V, respectively. For autoMS/MS, the MS1 scan rate was 2 Hz and the MS/MS scan rate was 0.5 Hz. A collisional energy formula was used with a slope of 2.0 and offset of 0. The max precursors per cycle was set to 2.

### Data Analysis

Mass spectra of intact species and dissociated subunits were analyzed using UniDec v6.0.3^52^ and compared with known interactions reported on UniProt. Deconvolution, isotopic fitting, and matching of fragment ions for both native and denaturing experiments were performed in MASH Native v1.1.^53^ To determine the number of detected native proteoforms, the MS1 scans were extracted from one SEC-IEC-MS experiment, split into one-minute windows, and deconvoluted in UniDec. Total ion chromatograms in Figure 1 are blank subtracted. Peak capacity in 2D-LC was estimated using the Davis equation,^38,39^ with ^1^D SEC peak capacity calculated according to the method of Hagel^54^ and ^2^D IEC peak capacity calculated according to the method of Neue for gradient elution.^32^

## Acknowledgements

The authors thank Dr. Christopher J. Wike (PolyLC) for providing columns and advice essential to this work. The authors also thank Dr. Rebecca Glaskin, John Sausen, and Jonathan Osborn (Agilent Technologies) for helpful discussions and instrument support. We would like to express sincere gratitude to the donor and their loved ones for their generous tissue donation. We thank James Anderson and Erin Halpin at the University of Wisconsin Organ and Tissue Donation for coordination of donor heart collection as well as Kalina Rossler and Scott Price for helping with the heart dissection. We would like to acknowledge the NIH R01 GM117058 (to Y.G. and S.J.), R01 HL109810, and S10 OD018475 (to Y.G.). E.A.C. would like to acknowledge support from the NIH Chemistry-Biology Interface Training Program NIH T32GM152341. H.T.R acknowledges the support of the National Heart, Lung, and Blood Institute of the National Institutes of Health under Award Number T32HL007936 through the UW-Madison Cardiovascular Research Center.

