## Supplemental Information for "Native Top-Down Proteomics of Endogenous Protein Complexes Enabled by Online Two-Dimensional Liquid Chromatography"

### Table of Contents

Supplementary Tables

Table S1. List of standard proteins used for method development…..………………………….…3

Table S2. Source and collision cell energies used for native 2D-LC-MS/MS experiments……3

Supplementary Figures

Figure S1. Effect of undersampling on peak capacity in 2D-LC…..……………………………...4

Figure S2. Solvent mismatch in online SEC-IEC……………….……………………………...4

Figure S3. Schematic representation of active solvent modulation……………..…………………..5

Figure S4. Development of SEC-IEC using standard proteins…..………………………………6

Figure S5. Design of Segmented Full-Window MS/MS method…...……………..……………...7

Figure S6. Representative LDHB and G3P fragment ions.. …..……………..…………..…...7

Figure S7. Detection of OXCT1 dimer, FH tetramer, and ECHM hexamer…………….............8

Figure S8. Representative OXCT1 fragment ions…….………………………………...............9

Figure S9. Representative ISOC1 fragment ions………………..……..……........................9

Figure S10. Native Top-Down MS analysis of endogenous human hemoglobin…...………...10

Figure S11. Reproducibility of SEC-IEC separation…………………………….…………......11

Figure S12. Comparison of native proteoforms detected by 1D and 2D-LC……...…..............12

Figure S13. Total ion chromatograms from 1D SEC and 1D IEC...............………….………...12

Figure S14. Additional complex-up analyses of human protein complexes…...…..………......13

Figure S15 Identification of CH60 by SEC-RPLC-MS/MS……………………….....................14

Figure S16. Detection of denatured subunits by SEC-RPLC-MS/MS…...………….………......15

Figure S17. Representative fragment ions from SEC-RPLC-MS/MS……..………….............16

### Supplementary Tables

**Table S1. List of standard proteins used for method development.**

| Protein | Abbreviation | UniProt ID | Mass (kDa) |
|---|---|---|---|
| Carbonic anhydrase | CA | P00921 | 29 |
| Enolase | Eno | P00924 | 93 |
| Ovalbumin | Ova | P01012 | 44 |
| Cytochrome C | Cyt | P00004 | 12 |
| Lysozyme | Lys | P00698 | 14 |
| Alcohol dehydrogenase | ADH | P00330 | 147 |
| Trypsinogen | Trp | Q29463 | 24 |

**Table S2. Source and collision cell energies used for native 2D-LC-MS experiments.** MS conditions used for the Agilent 6545XT Q-TOF.

| | | "MS1" | | | "MS2" | | |
|---|---|---|---|---|---|---|---|
| Injection # | SEC Cut # | Fragmentor (V) | Skimmer (V) | Collision Energy (V) | Fragmentor (V) | Skimmer (V) | Collision Energy (V) |
| 1 | 1 | 300 | 220 | 40 | 300 | 220 | 160 |
| | 2 | 250 | 150 | 20 | 250 | 150 | 120 |
| | 3 | 250 | 150 | 20 | 250 | 150 | 80 |
| | 4 | 250 | 100 | 0 | 250 | 100 | 40 |
| 2 | 1 | 300 | 220 | 40 | 300 | 220 | 100 |
| | 2 | 250 | 150 | 40 | 250 | 150 | 80 |
| | 3 | 250 | 150 | 20 | 250 | 150 | 60 |
| | 4 | 250 | 100 | 0 | 250 | 100 | 40 |
| 3 | 1 | 300 | 220 | 40 | 300 | 220 | 180 |
| | 2 | 250 | 150 | 40 | 250 | 150 | 160 |
| | 3 | 250 | 150 | 20 | 250 | 150 | 100 |
| | 4 | 250 | 100 | 0 | 250 | 100 | 60 |

### Supplementary Figures

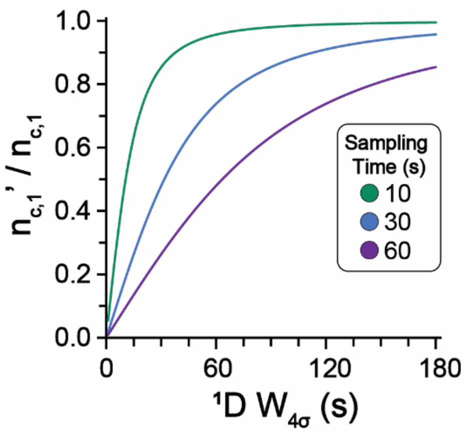


**Figure S1. Effect of undersampling on peak capacity in 2D-LC.** The loss of peak capacity due to undersampling is represented as the ratio of the Davis-corrected $^{1}$D peak capacity ($n'_{c,1}$) and the original $^{1}$D peak capacity ($n_{c,1}$). This result is plotted against $^{1}$D baseline peak width ($W_{4\sigma}$) for various sampling times, or frequency of consecutive fraction collection.

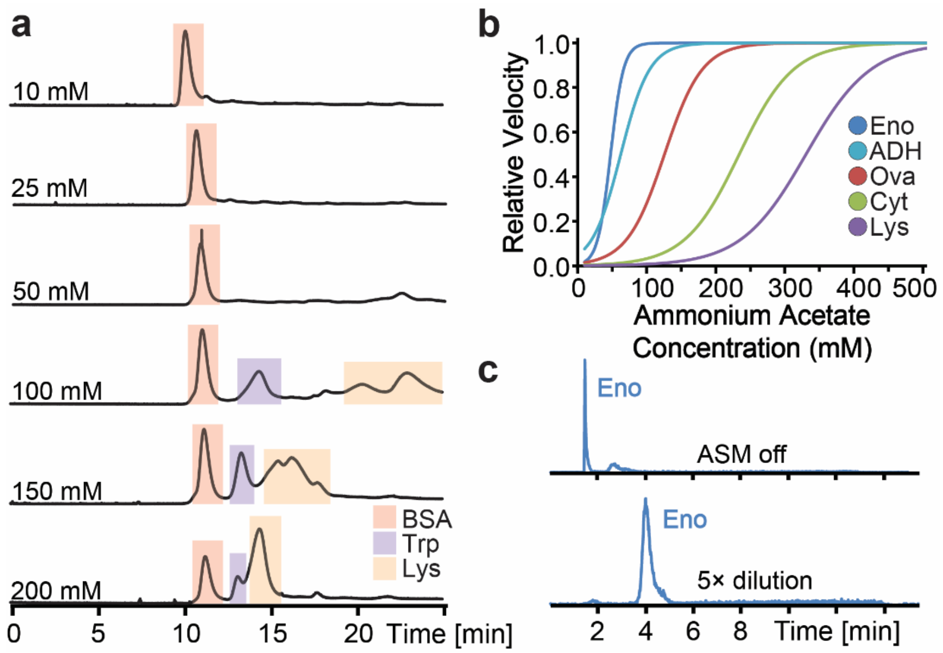


**Figure S2. Solvent mismatch in online SEC-IEC. a)** Effect of mobile phase ammonium acetate concentration on the SEC separation of three standard proteins. **b)** Plot of retention behavior for standard protein in mixed-bed IEC. Relative velocity is equal to the velocity of the solute divided by the velocity of the mobile phase. Solute velocities are calculated at each mobile phase condition according to the LSS model. **c)** Effect of ASM on the retention of Eno when transferred from $^{1}$D SEC (200 mM ammonium acetate) to $^{2}$D IEC.

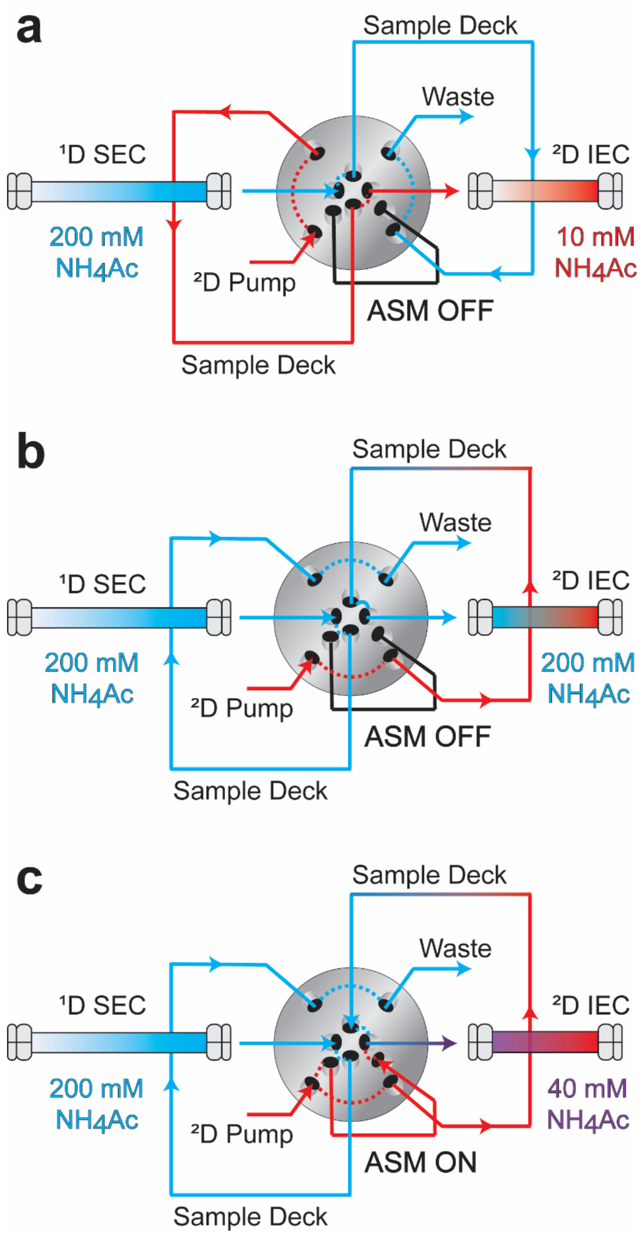


**Figure S3. Schematic representation of active solvent modulation. a)** In this scenario, no sample is being transferred from $^1$D SEC to $^2$D IEC, and the flow paths from each pump to its respective column are full of the proper solvent. **b)** Without ASM, switching the valve position results in a fraction of $^1$D mobile phase being injected into the 2D column, typically resulting in analyte breakthrough. **c)** With ASM, $^2$D mobile phase empties the sample loop and follows a parallel flow path through the ASM capillary directly to the head of the column, resulting in dilution of the transferred $^1$D effluent.

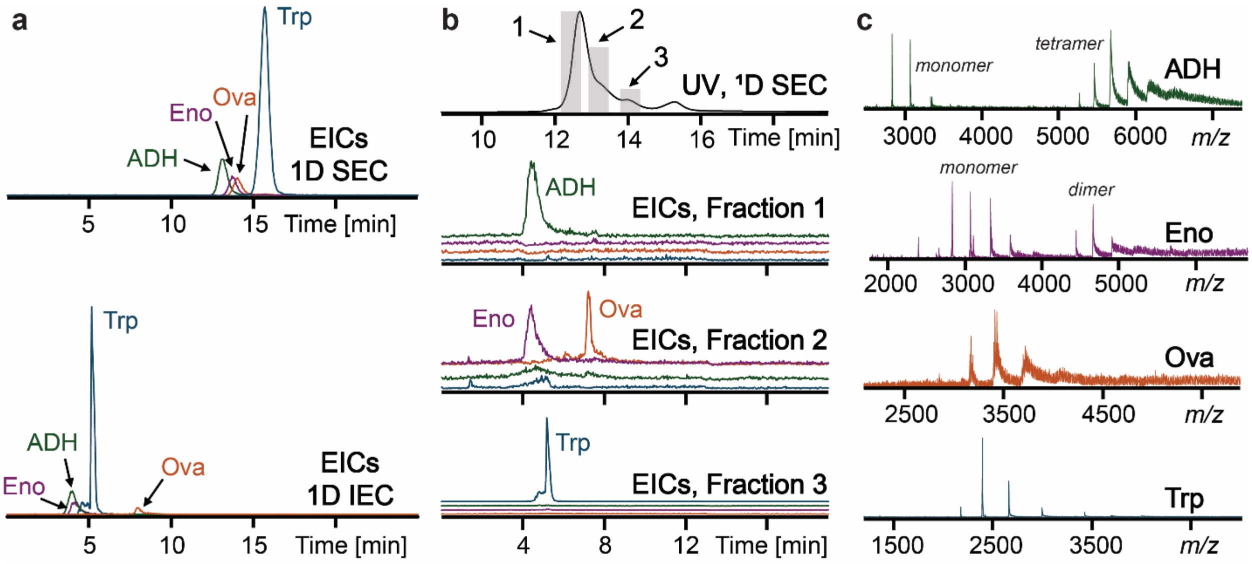


**Figure S4. Development of SEC-IEC using standard proteins. a)** Extracted ion chromatograms (EICs) from 1D separations of four standard proteins by SEC and IEC. **b)** Separation of standard proteins by SEC-IEC, including the UV trace from $^{1}$D SEC, location of three cuts (fractions), and the $^{2}$D IEC separation of each cut. **c)** Native mass spectra of standard proteins fully resolved by SEC-IEC.

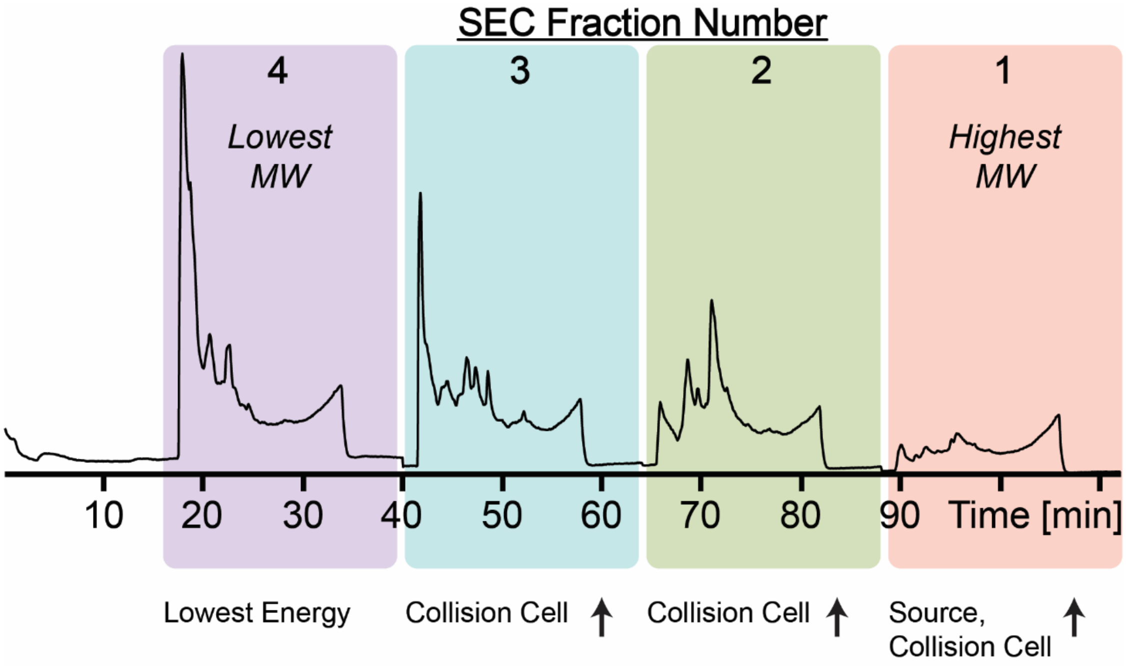


**Figure S5. Design of Segmented Full-Window MS/MS method.** Labeled total ion chromatogram from a 2D-LC-MS experiment. Source and collision cell energy was adjusted between SEC cuts to properly desolvate and dissociate protein complexes over a wide range of molecular weights and labilities. More detailed conditions are described in **Table S2**.

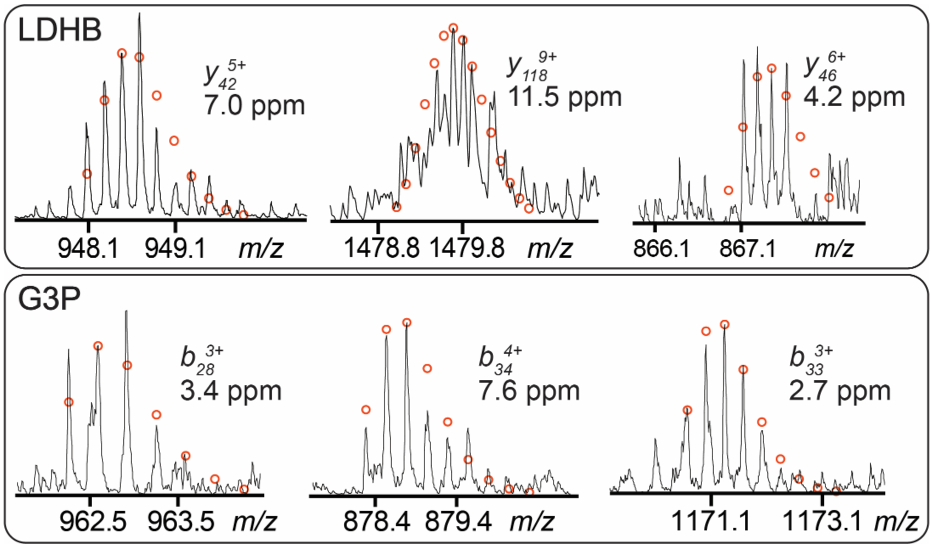


**Figure S6. Representative LDHB and G3P fragment ions.** Representative fragment ions from native top-down MS of L-lactate dehydrogenase B (LDHB) and glyceraldehyde 3-phosphate dehydrogenase (G3P).

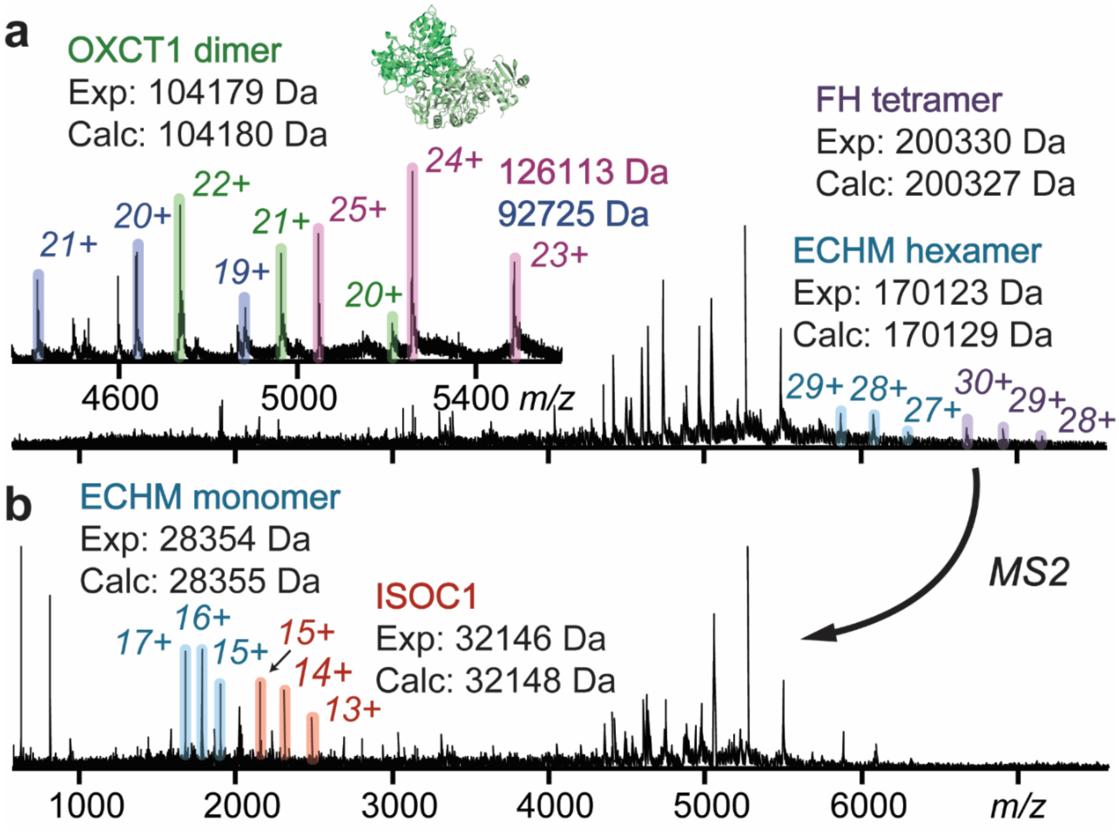


**Figure S7. Detection of OXCT1 dimer, FH tetramer, and ECHM hexamer. a)** Lower energy MS1 scan showing detection of the mitochondrial succinyl-CoA:3-ketoacid coenzyme A transferase 1 (OXCT1) dimer, the mitochondrial fumarate hydratase (FH) tetramer, and the mitochondrial enoyl-CoA hydratase (ECHM) hexamer. **b)** Higher energy full-window MS2 scan with detection of isochorismatase domain-containing protein 1 (ISOC1) and dissociated ECHM monomer.

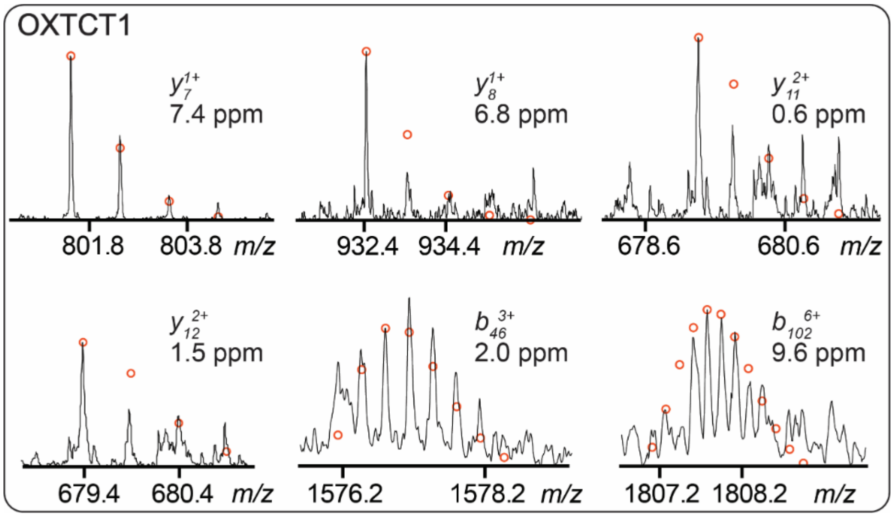

**Figure S8. Representative OXCT1 fragment ions.** Representative fragment ions from native top-down MS of mitochondrial succinyl-CoA:3-ketoacid coenzyme A transferase 1 (OXCT1).

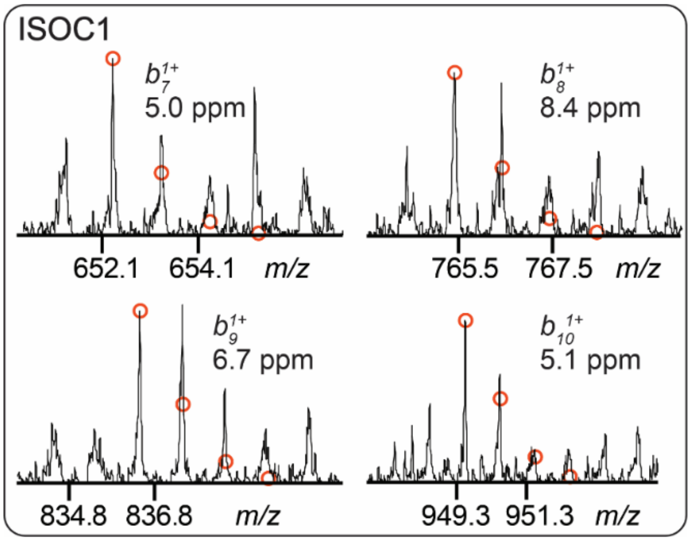

**Figure S9. Representative ISOC1 fragment ions.** Representative fragment ions from native top-down MS of isochorismatase domain-containing protein 1 (ISOC1).

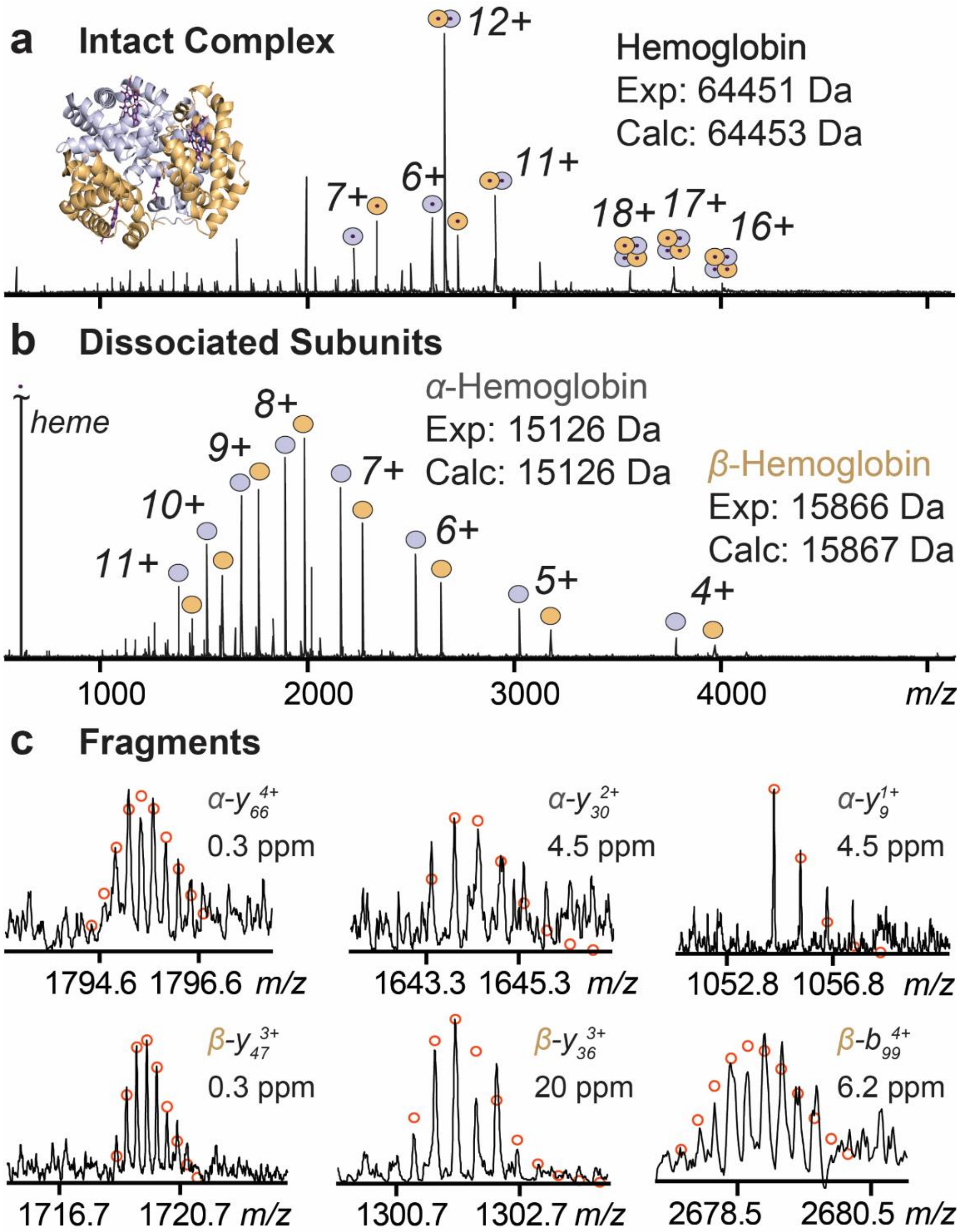


**Figure S10. Native Top-Down MS analysis of endogenous human hemoglobin. a)** Detection of the heme-bound heterotetramer, heterodimer, and monomers at the lowest energy condition tested. **b)** Dissociation of the subunits and bound heme groups upon increasing collisional energy. **c)** Representative fragment ions from both hemoglobin subunits with theoretical isotopic fits and high mass accuracy.

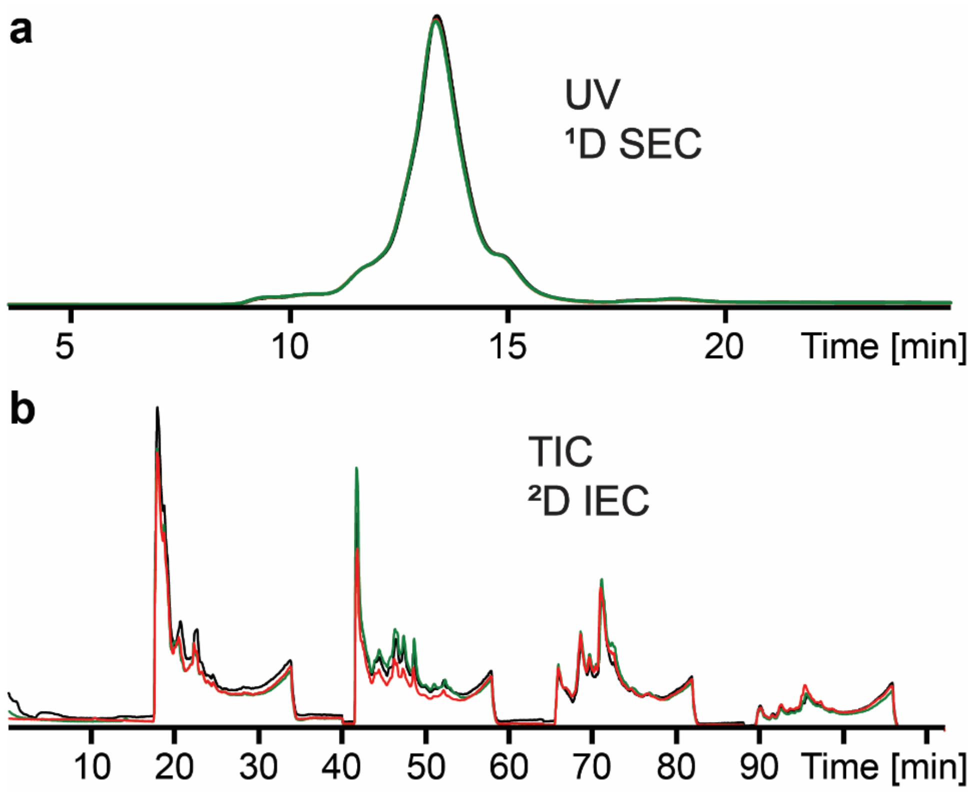


**Figure S11. Reproducibility of SEC-IEC separation.** Overlays of the $^{1}$D SEC UV trace and the total ion chromatogram (TIC) from three replicate injections.

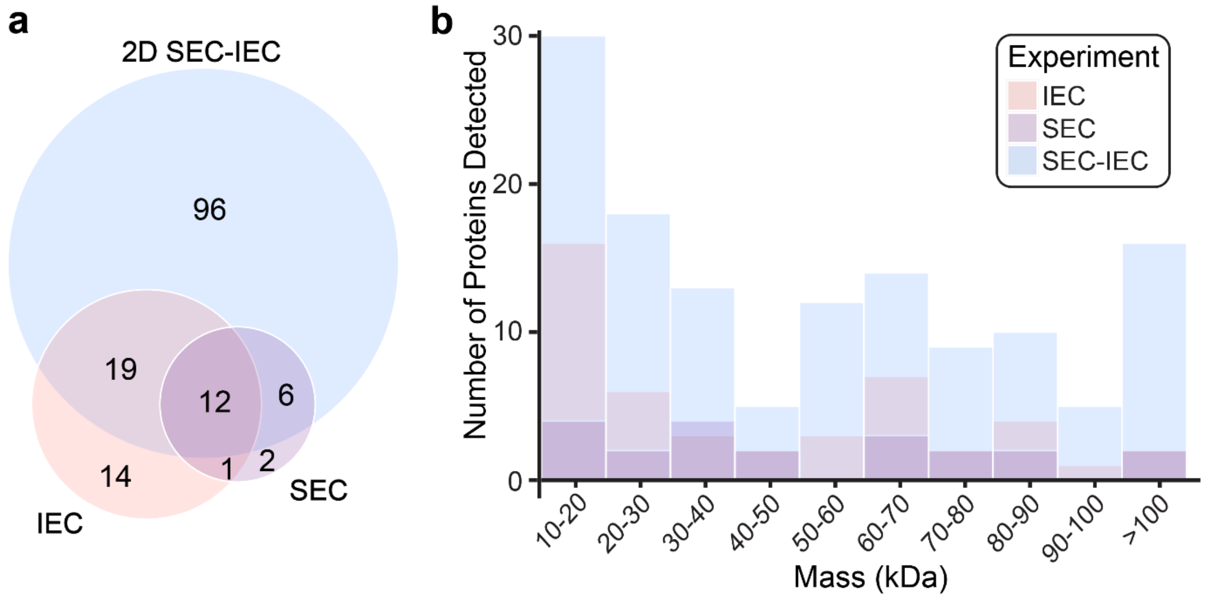


**Figure S12. Comparison of native proteoforms detected by 1D and 2D-LC. a)** Venn diagram showing unique and overlapping native proteoforms detected by 1D and 2D-LC methods. **b)** Histogram displaying the distribution of masses detected by 1D and 2D-LC methods.

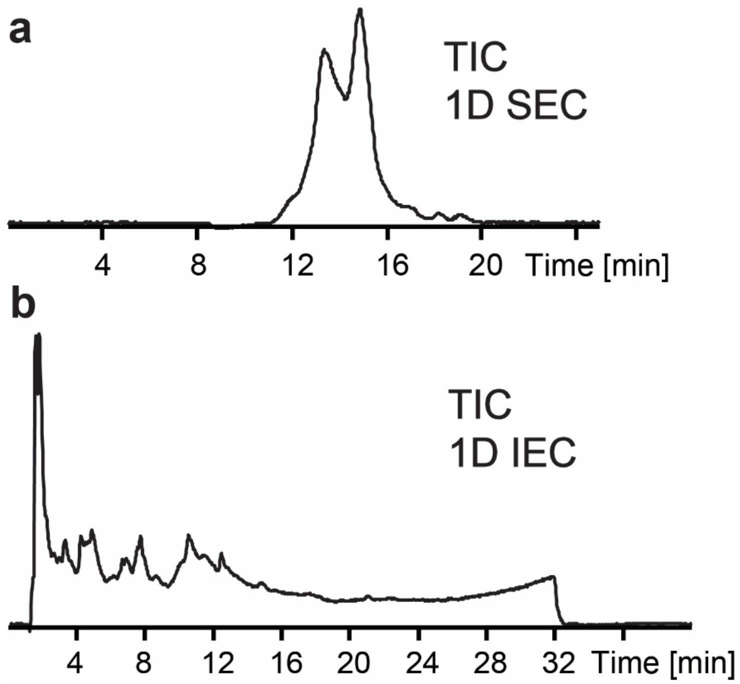


**Figure S13. Total ion chromatograms from 1D SEC and 1D IEC.**

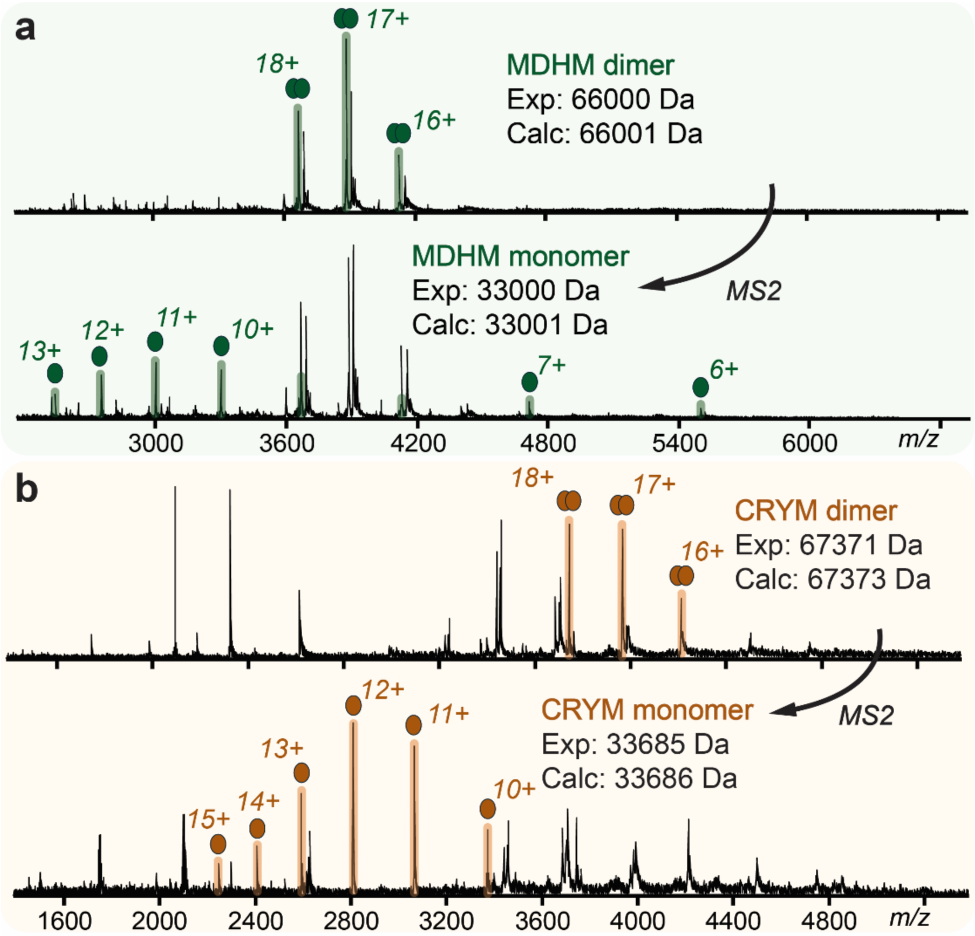


**Figure S14. Additional complex-up analyses of human protein complexes. a)** Detection of the intact complex and dissociated subunits of mitochondrial malate dehydrogenase (MDHM). **b)** Detection of the intact complex and dissociated subunits of ketimine reductase mu-crystallin (CRYM).

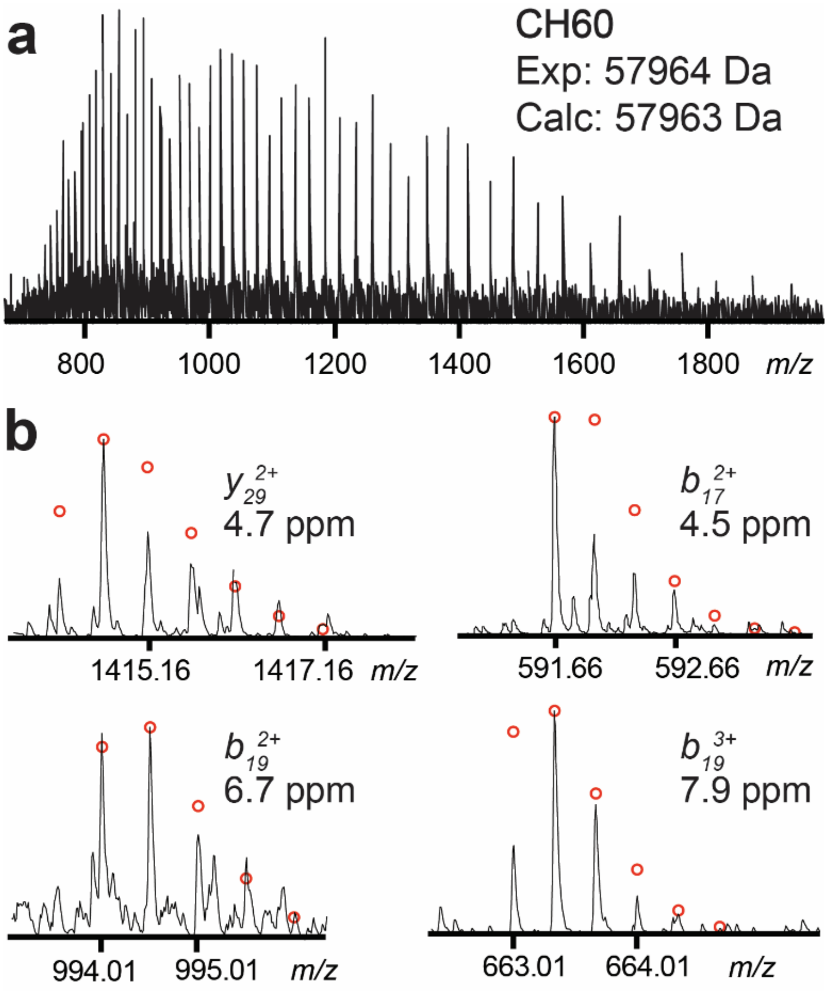


**Figure S15. Identification of CH60 by SEC-RPLC-MS/MS. a)** Mass spectra of CH60. The experimental and calculated masses are average masses. **b)** Representative CH60 fragment ions with theoretical isotopic fits and high mass accuracy.

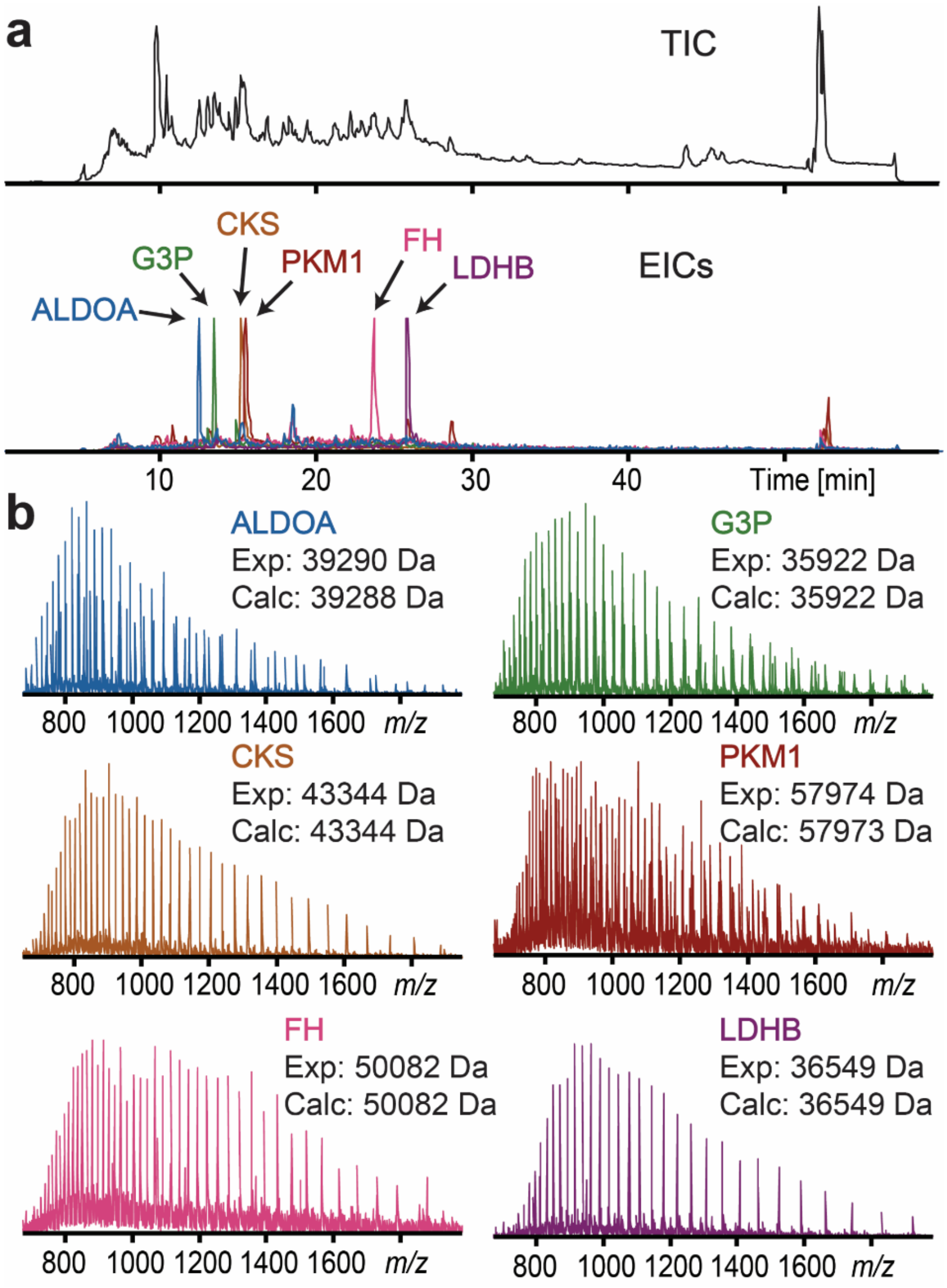


**Figure S16. Detection of denatured subunits by SEC-RPLC-MS/MS. a)** SEC-RPLC total ion chromatogram (TIC) and representative extracted ion chromatograms (EICs) from various proteins previously detected by SEC-IEC. **b)** Mass spectra corresponding to EICs. Experimental and calculated masses are average masses.

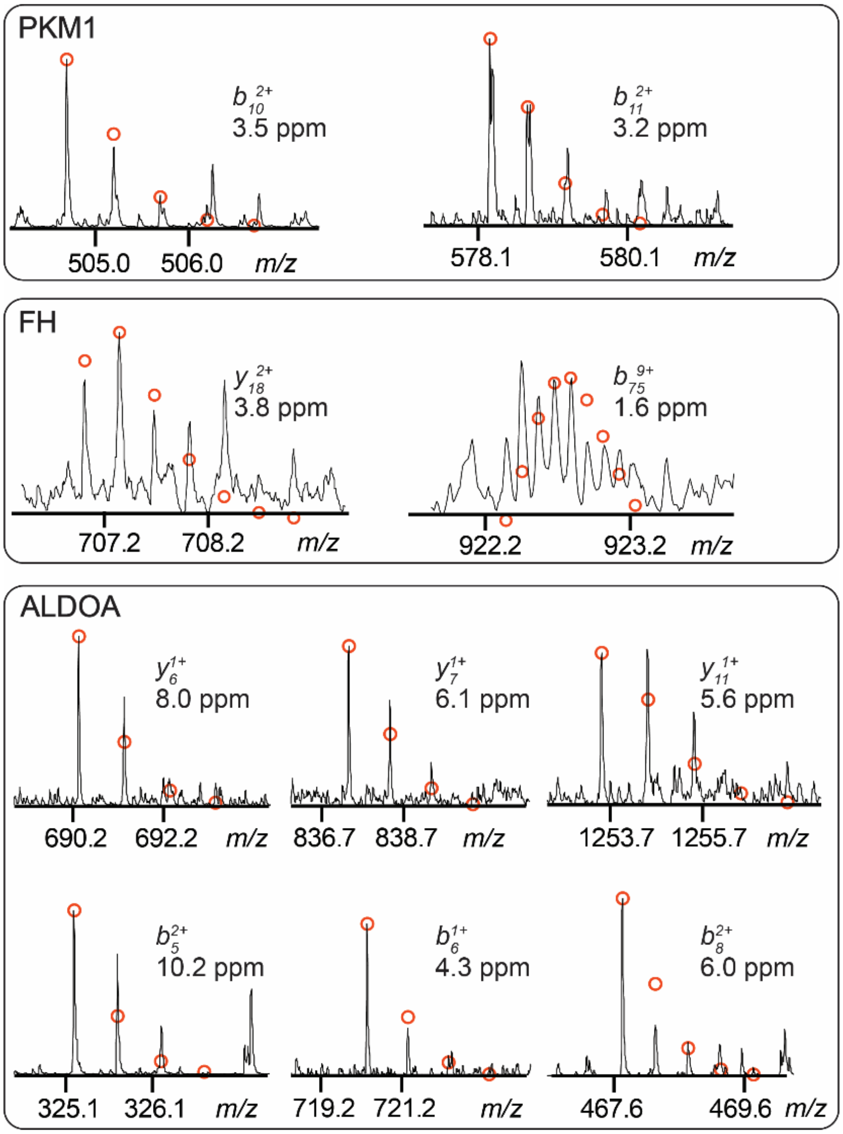


**Figure S17. Representative fragment ions from SEC-RPLC-MS/MS.** Representative fragment ions from SEC-RPLC confirming the identities of species inconclusively identified by SEC-IEC.
